# Evidence for co-evolution of masking and circadian phase in *Drosophila melanogaster*

**DOI:** 10.1101/2020.11.24.395558

**Authors:** Arijit Ghosh, Pragya Sharma, Shephali Dansana, Vasu Sheeba

## Abstract

Heritable variation in the timing or circadian phases of rhythmic events with respect to daily time cues gives rise to chronotypes. Despite its importance, the mechanisms (clock or non-clock) regulating chronotypes remain elusive. Using artificial laboratory selection for divergent phasing of emergence of adults from pupae, our group has derived populations of *Drosophila melanogaster* which are *early* and *late* chronotypes for eclosion rhythm. Several circadian rhythm characteristics of these populations have since been described. We hypothesized that our selection protocol has inadvertently resulted in selection for masking, a non-clock phenomenon, in the *early* chronotype due to the placement of our selection window (which includes the lights-ON transition). Based on theoretical predictions and previous studies on our populations, we designed experiments to discriminate between enhanced masking to light versus circadian clock mediated changes in determining enhanced emergence in the morning window in our *early* chronotypes. Using a series of phase-shift protocols, LD-DD transition, and *T*-cycle experiments, we find that our *early* chronotypes have evolved positive masking, and their apparent entrained phases are largely contributed by masking. Through skeleton *T*-cycle experiments, we find that in addition to the evolution of greater masking, our *early* chronotypes have also evolved advanced phase of entrainment. Furthermore, our study systematically outlines experimental approaches to examine relative contributions of clock versus non-clock control of an entrained behavior. Although it has previously been suggested that masking may confer an adaptive advantage to organisms, here we provide experimental evidence for the evolution of masking as a mean of phasing of an entrained rhythm that can complement clock control of an entrained behavior.

## Introduction

Circadian clocks adaptively schedule behavior and physiology to occur at a specific time of the day. Such scheduling is believed to be critical for maintaining our general health and well-being (Roenneberg et al., 2012; Vaze et al., 2014). Heritable variations in the timing (or phasing) of rhythmic events with respect to daily time cues result in what is referred to as chronotype variation (Infante-Rivard et al., 1989; Roenneberg et al., 2007). A clear example is that of variation in midsleep timing among humans on free days. While most humans will fall in the category of ‘normal’ or ‘neither’ chronotype, some individuals tend to fall asleep relatively early in the evening and wake up early in the mornings, hence referred to as *‘early’* chronotypes or *‘larks’* while there are those among us who prefer very late sleep timings and associated late wake timings, also referred to as *‘late’* chronotypes or *‘owls’* (Randler et al., 2017). Chronotypes are primarily controlled by the circadian clocks. Studies, including those on humans, have suggested that variation in entrained phases arise due to differences in underlying clock properties such as length of the intrinsic period of circadian clocks, phase/ velocity response curves (PRCs/ VRCs), amplitude of the circadian clock, inter-oscillator coupling, and amplitude of the zeitgeber (Aschoff and Pohl, 1978; Bordyugov et al., 2015; Daan and Pittendrigh, 1976; Granada et al., 2013; Johnson et al., 2003; Roenneberg et al., 2012; Swade, 1969).

While light can bring about a change in phase of an entrained rhythm by influencing the phase of circadian clock (Saunders et al., 1994; Schlichting and Helfrich-Förster, 2015), one cannot rule out aspects of direct effects of light. Such exogenous environmental influences on endogenously generated circadian rhythms which obscure aspects of circadian clock expression is referred to as masking (Aschoff, 1960a; Fry, 1947; Mrosovsky, 1999). Traditionally, non-involvement of the circadian clock has been considered an essential criterion defining masking, however, Terry Page (Page, 1989) and Nicholas Mrosovsky (Mrosovsky, 1999) make a strong case for the importance of masking as a complement to circadian clock regulation of daily rhythms. While there are studies which suggest that masking responses are present in animals without a functional clock (clock mutants or surgical ablation of the central clock) (Redlin and Mrosovsky, 1999; Wheeler et al., 1993), Aschoff had argued that there might be time-of-day dependence in certain masking responses (eliciting activity in blind male hamsters in the presence of mates) (Aschoff and Honma, 1999; Aschoff and von Goetz, 1988b). Positive masking refers to the masking response which elicits the beginning of a behavior, while negative masking refers to the inhibition or ceasing of the behavior. Traditionally studied in mammals with respect to locomotor activity rhythms, masking has also received some attention in insects, in both locomotor activity and eclosion rhythms (Hamblen-Coyle et al., 1992; Kempinger et al., 2009; Lu et al., 2008; Rieger et al., 2003; Sheppard et al., 2015; Thakurdas et al., 2009; Wheeler et al., 1993).

The act of emergence of a pharate adult fly from its pupal case or eclosion is developmentally gated, and a population level rhythm. It was also among the earliest circadian rhythms to be studied systematically (Chandrashekaran, 1967; Chandrashekaran and Loher, 1969; Engelmann, 1969; Harker, 1965a, 1965b; Pavlidis, 1967; Pittendrigh, 1954, 1967; Pittendrigh et al., 1958; Skopik and Pittendrigh, 1967; Zimmerman et al., 1968). The anatomical and physiological processes underlying eclosion have also been extensively investigated (Johnson and Milner, 1987; Krüger et al., 2015; Peabody and White, 2013; Selcho et al., 2017; Thummel, 2001). It is deemed to be amongst one of the most critical events in the lifetime of a holometabolous insect (Mcmahon and Hayward, 2016; Zitnan and Adams, 2005). The *Drosophila* eclosion rhythm has been shown to be regulated by the circadian pacemakers previously implicated in activity rhythms in addition to prothoracic gland (PG) clocks (Morioka et al., 2012; Myers et al., 2003; Pittendrigh and Bruce, 1959; Selcho et al., 2017; Zimmerman et al., 1968). Additionally, in contrast to several behavioral rhythms, the act of eclosion is free from the influence of motivational state, other behaviors, or interactions among individuals, while being susceptible to disruption when mutations are introduced in core clock genes like *period* (*per*) or *timeless* (*tim*) (Qiu and Hardin, 1996; Ruf et al., 2019; Sehgal et al., 1994) both under constant and cycling conditions. Thus, relative to other rhythms, it appears to be more sensitive to perturbations in the core clock. Eclosion rhythms, their entrainment to light/ dark cycles, temperature cycles, synergistic light and temperature cycles as well as molecular mechanisms of their entrainment have been studied in great detail in Drosophilid species (Emery et al., 1997; Kumar et al., 2006, 2007; Morioka et al., 2012; Myers et al., 2003; Nikhil et al., 2014, 2015; Nikhil, Abhilash, et al., 2016; Pittendrigh, 1966; Pittendrigh and Bruce, 1959; Pittendrigh and Minis, 1972; Prabhakaran et al., 2013; Qiu and Hardin, 1996; Vaze and Sharma, 2013). The rhythm in eclosion is modulated by the lights-ON signal (Chandrashekaran and Loher, 1969; Engelmann, 1969; Pittendrigh, 1967; Thakurdas et al., 2009), even though wing-expansion, the last behavioral event of the *Drosophila* adult eclosion sequence (Fraekkel, 1935), is not affected by the same (McNabb and Truman, 2008), suggesting that the lights-ON signal may have a role specific to the act of eclosion itself. It was suggested previously, that the lights-ON signal may have two distinct effects on the timing of eclosion of flies - a) stimulation of eclosion hormone (EH) release, and b) reduction in the latency of eclosion relative to EH release (Baker et al., 1999; McNabb and Truman, 2008).

With reference to *Drosophila melanogaster*, Hamblen-Coyle and colleagues first reported a lights-ON peak in locomotor activity rhythm which they designated as a ‘ *startle*’ effect. It was seen that even though flies carrying core circadian clock mutations adopt distinct phases in evening peak timings, their morning peak (lights-ON peak) phases were not very different (Hamblen-Coyle et al., 1992). Later it was shown that under LD12:12, even without a functional clock, animals exhibited this lights-ON peak, but it was absent when shifted to constant darkness (DD) (Wheeler et al., 1993). In case of locomotor activity rhythm, Rieger and colleagues showed that under laboratory LD12:12 conditions, the lights-ON peak and the morning peak are indistinguishable as they overlap with each other (Rieger et al., 2003). These two peaks can be separated from each other under different photoperiods, where the lights-ON peak is still phase-locked to the dark-to-light transition, but the morning peak is advanced or delayed under short or long photoperiod respectively (Rieger et al., 2003). Artificial moonlight can make fruit flies nocturnal, and this nocturnal light is known to induce strong locomotor activity in flies via masking (Kempinger et al., 2009). Recent detailed genetic dissections of masking in *Drosophila* have revealed complex pathways mediating light-induced masking of locomotor activity (Rieger et al., 2003). Furthermore, Lu and colleagues demonstrated a circadian rhythm in light-induced locomotor activity against a background of DD and showed that the circadian clock genes *timeless* and *clock* are involved in regulation of this masking response (Lu et al., 2008; Sheppard et al., 2015). Taken together, the above studies on effect of light on locomotor activity rhythms and eclosion suggest that masking can affect the timing of a circadian rhythm in *Drosophila*.

To investigate the immediate effects of light on the timing of eclosion rhythms, we use a set of four populations of *D. melanogaster* which could potentially have evolved a masking response due to the nature of the selection regime *(early_1-4_*- described below). These populations are part of an on-going long-term experimental evolution study at the Chronobiology laboratory, JNCASR. The primary goals of creating these populations were - a) to demonstrate the adaptive significance of phasing of circadian rhythm (here, eclosion rhythm) b) to then study the associated clock properties mediating phase divergence. Indeed our populations selected for morning emergence *(early_1-4_)* exhibit shorter free-running period (FRP) compared to those selected for evening emergence (*late_1-4_*) and the *control_1-4_* populations, which did not undergo any selection for timing of emergence (Kumar et al., 2007; Nikhil, Abhilash, et al., 2016). Over the years several studies from our group have shown that entrainment of these divergent chronotypes to light-dark (LD) cycles cannot be fully explained by exclusively invoking either the parametric or non-parametric models of entrainment (Abhilash and Sharma, 2020; Kumar et al., 2007; Vaze, Nikhil, et al., 2012). It was also hypothesized that differences in inter-oscillator (A / master / central oscillator and B / slave / peripheral oscillator) coupling might explain the chronotype divergence in eclosion rhythm (Abhilash et al., 2019; Nikhil, Abhilash, et al., 2016). At the time of writing, these populations have undergone nearly 340 generations of selection.

We reasoned that our selection lines may provide material to examine some aspects of masking and circadian control of a rhythmic phenomenon because the design of our selection regime is such that flies of the *early* populations have, over generations, been forced to emerge in a window around lights-ON (3 hours prior to lights-ON till 1-hour post lights-ON). It is possible that this protocol has inadvertently resulted in selection for the phenomenon of masking. Therefore, among other clock related factors that have been uncovered to have changed in these populations previously (Abhilash and Sharma, 2020; Nikhil et al., 2015; Nikhil, Vaze, et al., 2016; Vaze, Kannan, et al., 2012) we hypothesized that the phase divergence among *early, control* and *late* chronotypes are partly due to differences in masking - specifically, that *early* populations exhibit a high degree of masking. We designed several experiments to examine whether *early* flies exhibit enhanced positive masking in eclosion rhythm compared to *control*, and *late* flies. Our results demonstrate that - a) *early* chronotype flies have indeed evolved significantly more positive masking compared to *control* and *late* flies, b) under full photoperiod, apparent entrained phases of *early* flies are largely contributed by masking, c) under skeleton photoperiod, *early* flies do show phase lability, and retain advanced phase of entrainment compared to *control* and *late* flies to different *T*-cycles, suggesting that our selection indeed has selected for greater masking alongside selection for advanced phase of entrainment.

## Materials and methods

### Selection protocol and fly maintenance

F our sets of genetically independent *Drosophila melanogaster* populations were used to artificially select for morning eclosion *(early* populations) and evening eclosion *(late* populations) timing - henceforth, denoted as *early_(1-4)_*, *control_(1-4)_* (no selection imposed) and *late(1-4)*. At each generation, ~300 eggs are collected and placed in glass vials which are maintained in a light-proof, temperature-controlled cubicle in a light-dark cycle of 12/12 hours (LD12:12), 25±0.5 °C, and 65±5% RH. Flies emerging from ZT21-ZT01 (ZT0 is <u>z</u>eitgeber <u>T</u>ime 0, when light comes ON) are collected to form the breeding pool for the next generation of *early* flies, while flies emerging from ZT9-ZT13 form the breeding pool for the next generation of *late* flies. Flies emerging throughout the day are collected to make up the breeding pool for the next generation of the *control* flies. This collection goes on for 3-4 days and total ~1200 adult flies with ~1:1 sex ratio of each of the 12 populations are maintained in Plexiglas™ cages (25 cm×20 cm×15 cm) with petri plates with banana-jaggery culture media. The flies are maintained on a 21-days discrete generation cycle, and all experiments were done with progeny of flies that experienced one generation of common rearing (standardized) to avoid confounding factors due to maternal effects (Bonduriansky and Day, 2009). For more details of the selection regime, see Kumar *et al.* (Kumar et al., 2007), and Abhilash *et al.* (Abhilash et al., 2019). All experiments were done between generations 320 - 330.

### Behavioral experiments

Before each experiment, ~300 eggs were collected and placed in 10 vials each for all 12 standardized populations. After egg collection, the vials were maintained in different regimes specific to each experiment. Emerged flies were counted every two hours or half an hour after the assay started, depending on the experimental regime. Briefly, all rhythm assays in *Fig. 2 and 3* were carried out with half hour resolution 12 hours around lights-ON and in assays depicted in *Fig. 4 and 5* (full and skeleton T-cycle experiments), 2 hours resolution was used due to logistic constraints. To account for differences in development time, if any, fly counts from the first emergence cycle were excluded from analysis. All 12 populations were assayed in parallel. In all experiments, temperature (25±.5 °C), and light intensity (~70 lux, from a white LED source) were kept constant. Other details of the light regime are mentioned in each experiment separately, and light ON-OFF times (step wise) were programmed with a TM619 timer (Frontier Timer, Pune, India) in the incubators (DR-36VL, Percival Scientific, Perry, USA) in which the experiments were performed.

**Figure 1:**
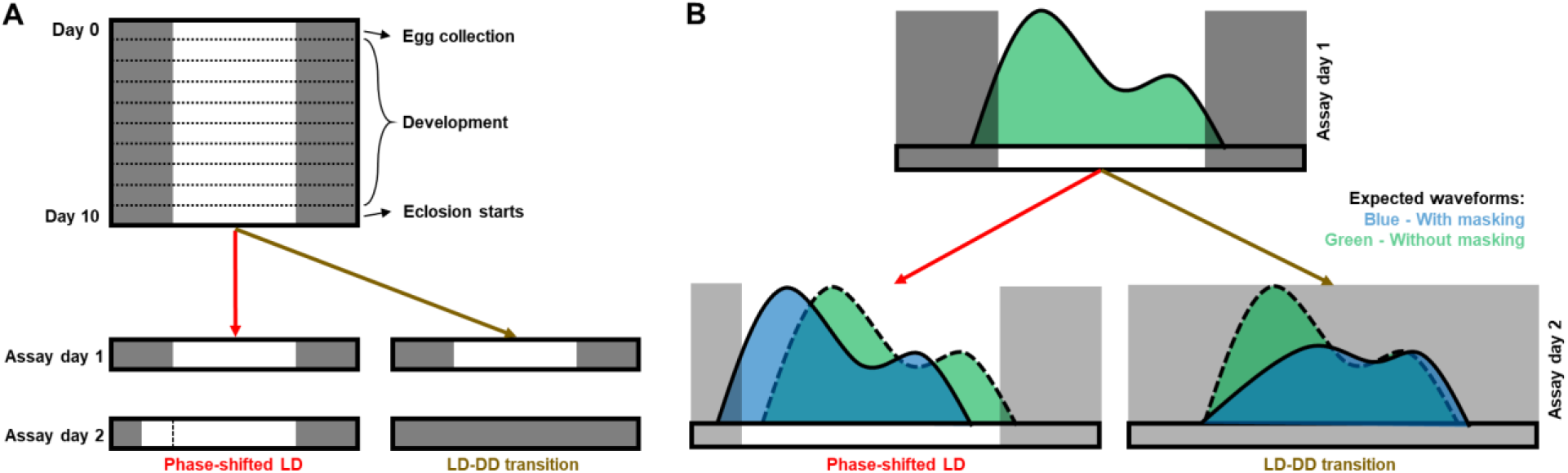
Schematics of experimental design and hypotheses. **A.** After egg collection, vials were kept under LD12:12 till 9^th^ day. Emergence assay was started on the 10^th^ day *(Assay day 1),* and flies were counted every two/half an hour interval depending on experiments and time points. On the 11^th^ day *(Assay day 2),* vials were placed under a 3-hour phase-advanced light schedule (Phase-shifted LD) or complete darkness (LD-DD transition). **B.** The schematic shows expected waveforms in case of complete circadian control (in green) or masking (blue). Dark rectangular shades depict duration of darkness.

**Figure 2:**
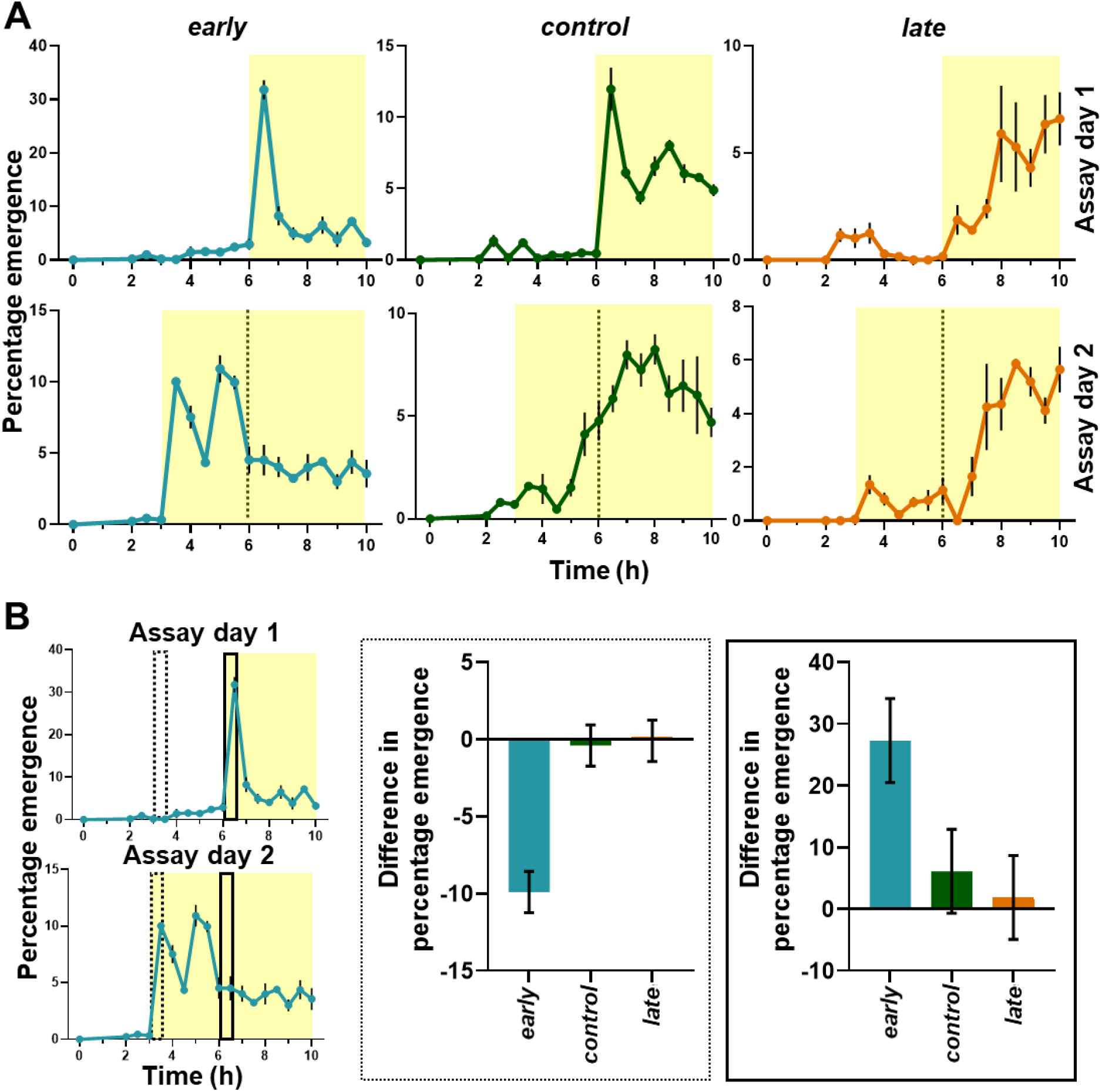
Emergence profile during first 10 hours on day 10 (post egg collection) in pre-shift *(Assay day 1)* and post-shift *(Assay day 2)* days and difference in percentage emergence between pre-shift *(Assay day 1)* and postshift *(Assay day 2)* days in two 0.5-hour time windows. **A.** *early* flies (blue line) advance their emergence waveform to emerge immediately after lights-ON on *Assay day 2.* The *control* (green line) and *late* (orange line) flies conserve their waveform on both days by not advancing their emergence in response to sudden advancement of the lights-ON stimulus. The yellow shading indicates the photophase in the LD cycle each day. Error bars are ± SEM. Grey dotted lines in the bottom panel indicates lights-ON time on previous day. **B.** The left panels show time windows used for analysis (dashed and solid rectangles respectively). In same time windows *early* flies show significantly higher (dashed rectangle region) or lower (solid rectangle region) emergence than that of *control* and *late* flies on *Assay day 2* than *Assay day 1,* showing this high emergence immediately after lights-on is specific to *early* flies.

### Analysis of data and statistics

Average profiles were constructed first by averaging over multiple cycles for a vial, and then by averaging over vials for each population. All statistical comparisons were made using 2-way or 3-way randomized block design ANOVA with selection regime and *T*-cycle (wherever applicable) as fixed factors and blocks (replicates) as the random factor. Results were deemed significant for main effect or interaction as applicable at α < 0.05. Post-hoc comparisons were carried out by a Tukey’s Honest Significant Difference (HSD) test. All statistics were performed in Statistica 7 (StatSoft, Tulsa, USA). Standard errors of means have been plotted as error bars in average profiles for ease of visualization. All 95% confidence intervals from the Tukey’s HSD are plotted in quantitative comparisons for visual hypothesis testing. Basic data processing and calculations were done with Microsoft Excel 365, and all graphs were plotted with Graphpad Prism 8. Criteria for “apparent entrainment” was T_observed_ must equal to T_environment_ (T_observed_ is the observed period of the eclosion rhythm and T_environment_ is the duration of the light/ dark cycle). T_observed_ was calculated with JTK-cycle (Hughes et al., 2010) employed in MetaCycle2d (Wu et al., 2016) with percentage eclosion data for each vial. All calculated phases are essentially phase relationships with the lights-ON signal. Peak phase is denoted by Ψ_Peak_ and measured in hours, and phase of Centre of Mass is denoted by Ψ_CoM_ and measured in degrees. Ψ_Peak_ was calculated as the time where maximum number of flies emerged in each vial and Ψ_CoM_ was calculated as a measurement of mean phase of emergence in polar coordinates for each vial (corrected for different lengths of *T*-cycles). Consolidation of emergence/ normalized amplitude (R) was also calculated in a polar coordinate system for eclosion data averaged over cycles, details of computation and usefulness of which can be found in a recent publication from our lab (Abhilash et al., 2019).

## Results

### Lights-ON elicits an immediate response in the *early* chronotypes

Previously, all eclosion rhythm assays on our populations were carried out with a maximum resolution of 2 hours and thus far no difference was detected in the peak phase (Ψ_Peak_) of *early* and *control* flies, both of which were found to emerge maximally at (ZT2) (Kumar et al., 2007; Nikhil, Abhilash, et al., 2016). To examine whether there are subtle changes in the onset or peak of emergence among stocks we increased the resolution to 0.5 hours for the first half of the day *(Assay day 1* and *2, Fig. 2A & 3A).* In fact, we now find that peak (Ψ_Peak_) emergence for both *early* (~30%) and *control* (~15%) flies occur at ZT0.5, i.e., immediately after lights-ON. Onset of emergence is similar as compared to 2-hours resolution assays done previously (Kumar et al., 2007; Nikhil, Abhilash, et al., 2016) across stocks.

**Figure 3:**
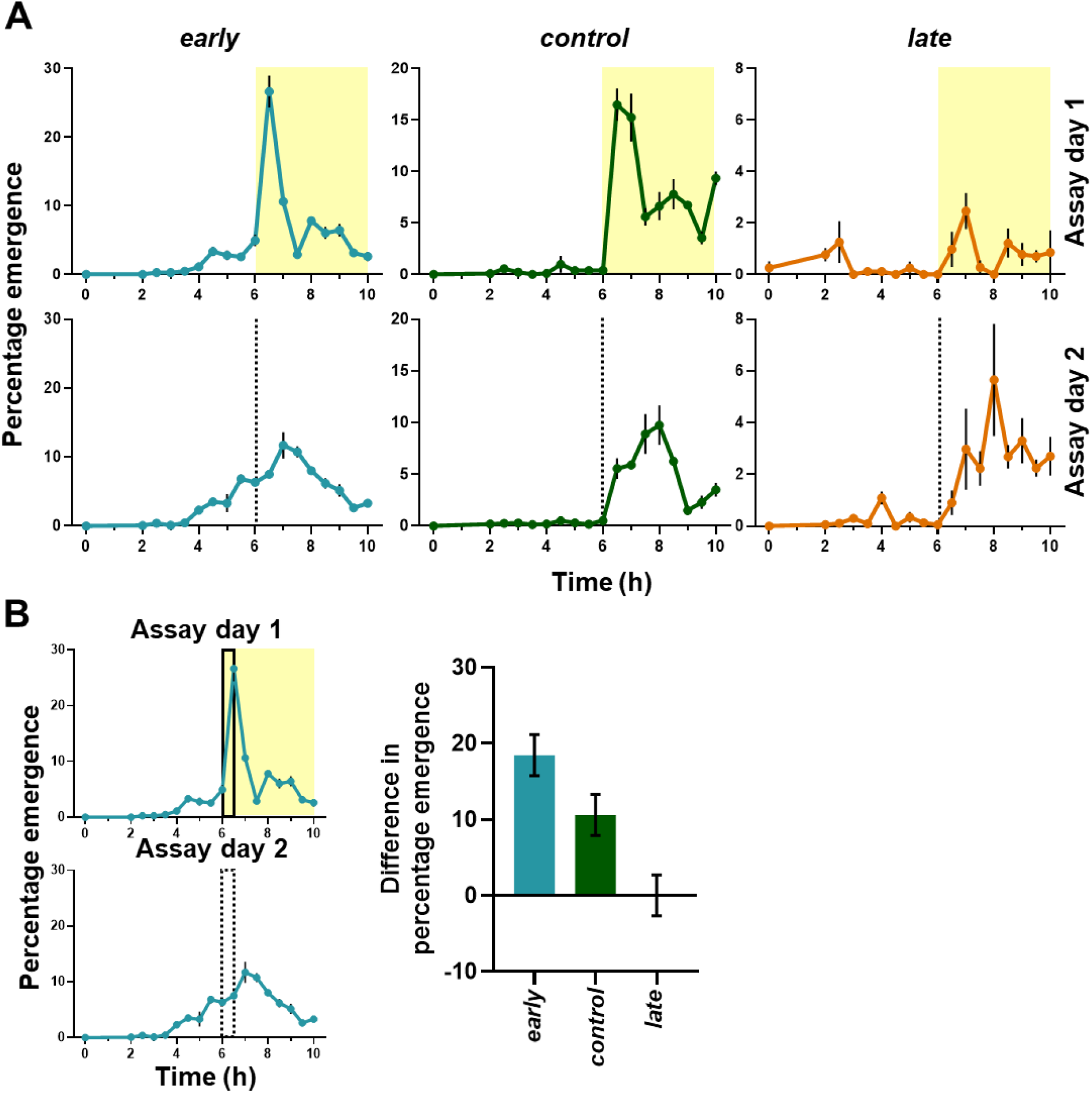
Emergence profile during first 10 hours on day 10 (post egg collection) (after 10 days) in LD *(Assay day 1)* and DD *(Assay day 2)* days and quantification of difference of percentage emergence between pre-shift and post-shift days from ZT0-0.5 (lights-ON phase followed from LD cycle). **A.** *early* flies attenuate their emergence to a great extent in absence of light stimulus in the earliest part of the subjective photophase, unlike their waveform in LD12:12. However, *control* and *late flies*, emerge with waveforms like under LD12:12, albeit with lower amplitude.. Error bars are ± SEM. Grey dotted lines in the bottom panel indicates lights-ON time on previous day. **B.** *early* flies show significantly higher difference than *control* and *late* flies. This suggests the high emergence of *early* flies immediately after lights-on was a masking response rather than a clock mediated response.

When lights-ON was advanced by 3 hours on the following day *(Assay day 2, Fig. 2A*), *early* flies exhibited high emergence in the first 0.5 hour, whereas the *control* and *late* flies emerged at times similar to that of the previous cycle *(Assay day 1, Fig. 2A).* Peak emergence of *early* flies occurred at ZT2 on *Assay day 2*. On both assay days, majority of *early* flies emerged immediately after lights-ON *(Fig. 2A).* The phase of peak emergence for *control* flies remained similar to *Assay day 1,* even after the light phase was advanced on *Assay day 2 (Fig. 2A*). Emergence waveform of *late* flies did not change from *Assay day 1* to *Assay day 2 (Fig. 2A*).

Based on the emergence profile of the *early* populations we quantified the difference in emergence during two specific time windows on the two experimental days. The first window (*solid rectangle, Fig. 2B)* depicts the time window of maximum emergence for *early* flies on *Assay day 1* (LD) and the second window *(dashed rectangle, Fig. 2B)* depicts the time window for maximum emergence (immediately after lights-ON) for *early* flies on *Assay day 2* (phase-shifted LD). If *early* flies show more masking than circadian control of their emergence, one would expect a large difference in levels of eclosion between the two days because they are strongly modifying their waveforms in response to lights-ON. We observe that the difference in percentage emergence during the first window (between *Assay days 1* and *2*) is significantly higher for *early* flies than for *control* and *late* flies, the latter two showing almost no change across days (*solid rectangle, Fig. 2B; F*_2,6_ = 200.3084, *p* < 0.05; *Supplementary table S1).* Similarly, in the second window, *early* flies show significantly higher change across Assay days 1 and 2 compared to *control* and *late* populations *(dashed rectangle, Fig. 2B; F*_2,6_ = 46.2010, *p* < 0.05; *Supplementary table S2).* This suggests that on *Assay day 2*, the circadian response of *early* flies is overridden by the immediate lights-ON response, thus exhibiting higher positive masking response compared to *control* and *late* flies. When considering two larger windows each of 2.5 hours (around lights-ON of pre-shift day) similar differences prevail *(Supplementary fig. S14),* showing that high emergence immediately after lights-ON is specific to *early* flies.

### *early* flies attenuate and delay their emergence under DD

Since the above experiments suggested that masking plays a prominent role in the timing of eclosion of *early* flies we then attempted to parse the relative contribution of circadian clock control on the emergence profile of *early* flies. Therefore, one set of cultures were shifted to constant darkness (DD - *Assay day 2, Fig. 1A right panels; LD-DD transition).* If the circadian clock has a strong control over the eclosion rhythm, the *early* flies should show high emergence in the early part of the subjective photophase in DD, which was not observed. There is a stark difference in emergence between two consecutive cycles during the 2-hour time window immediately after lights-ON which is much lower in *control* and almost nonexistent in *late* flies *(Fig. 3A and 3B right panel).* The sharp morning peak, an identifiable marker for our *early* flies was absent in DD (*bottom left panel, Fig. 3A*).

We quantified the difference in percentage emergence from ZT0-0.5 of first day (LD, *Assay day 1)* and the same phase in second day (DD, *Assay day 2*), and find that *early* flies show significantly larger difference than *control* and *late* flies, suggesting attenuated emergence immediately after starting of subjective photophase in DD *(Fig. 3B; F*_2,6_ = 55.4322, *p* < 0.05; *Supplementary table S3*). While a reduction in amplitude is expected under DD, it is significantly greater for *early* flies than *control* flies. This suggests the high emergence of *early* flies immediately after lights-ON is largely contributed by a masking response and less of a clock-controlled phenomenon.

### *early* flies consistently emerge close to lights-ON under both short and long *T*-cycles

*T*-cycles shorter than the FRP of the organism are expected to delay phases of circadian rhythms (phase relationships with zeitgeber) while *T*-cycles longer than the FRP of the organism advance phases of the circadian rhythms (Aschoff, 1965; Wheeler et al., 1993; Yadav et al., 2015). Therefore, we subjected our populations to a series of short (T20; LD10:10 and T22; LD11:11) and long (T28; LD14:14 and T26; LD13:13) *T*-cycles. We expected that eclosion rhythm of *control* and *late* flies, owing to strong control by circadian clocks will shift their phases in the predicted directions. The previous experiments *(Fig. 2 and 3*) suggested that positive masking of light governs the phase of eclosion rhythm of *early* flies to a large extent. We, therefore, hypothesized that the *early* flies will not modify phases of the eclosion rhythm in the predicted directions under short or long *T*-cycles - delay under T>24 hours and advance under T<24 hours.

Under T22, *early* flies maintain similar waveform as under T24 *(left panel top row, Fig. 4A).* The *control* flies showed clear phase delay *(middle panel top row, Fig. 4A),* whereas *late* flies showed even greater phase delay in response to a T22 cycle *(right panel top row, Fig. 4A)* compared to their waveforms in T24 cycles.

**Figure 4:**
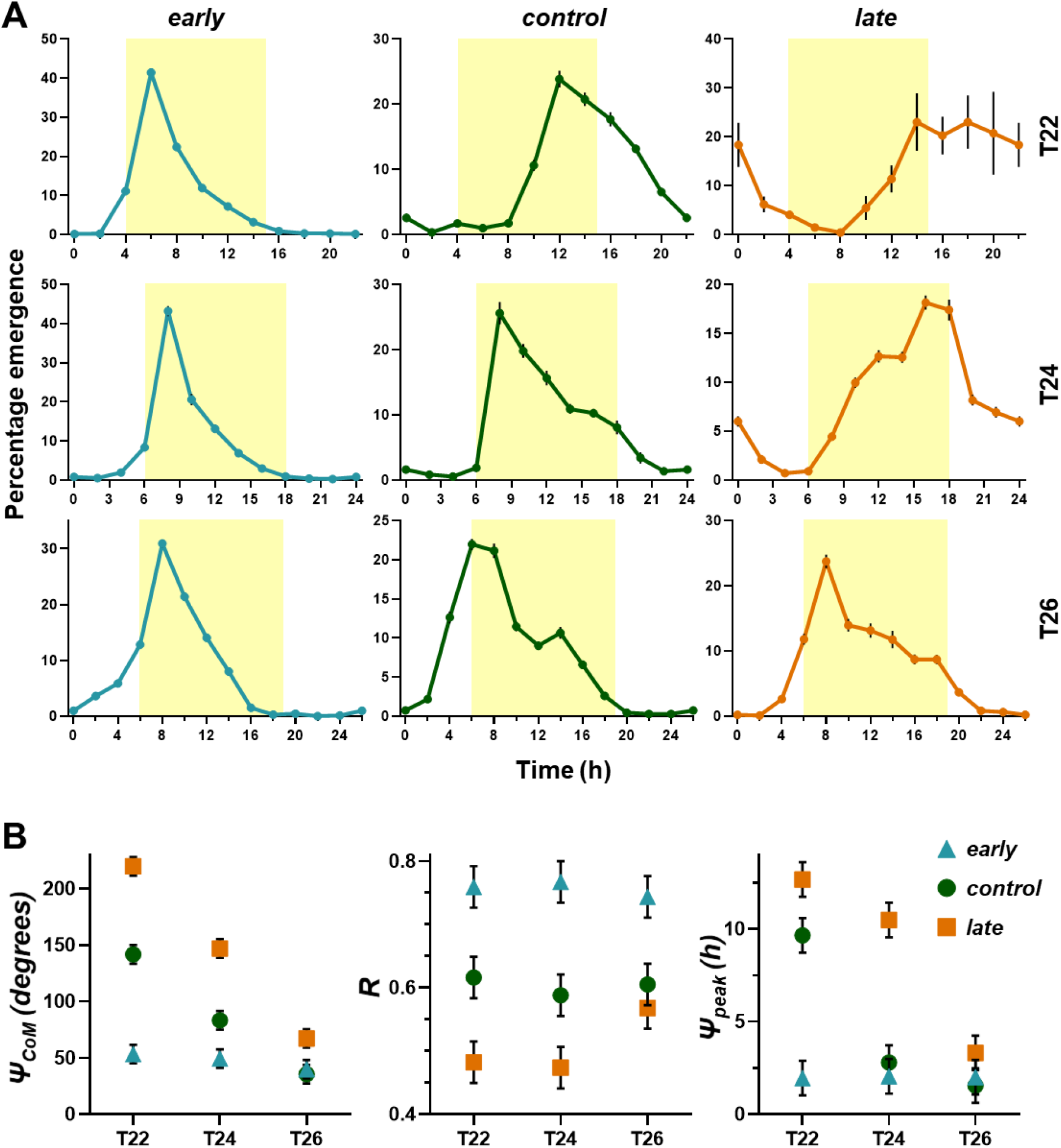
Emergence profile of *early, control,* and *late* flies under T22, T24, and T26 cycles and eclosion rhythm parameters under *T*-cycles. **A.** *early* flies do not change their emergence profile depending on *T*-cycles, whereas *control* and *late* flies shift their emergence profile in expected directions (phase advance in longer than 24-hour cycle and phase delay in shorter than 24-hour cycle). Yellow shading depicts the photophases of respective *T*-cycles. Error bars are ± SEM. **B.** Left panel: Centre of Mass (Ψ_CoM_ - in degrees), middle panel: Consolidation of emergence (R), right panel: Peak in ZT (Ψ_Peak_, in hours). Error bars are ±95% CI. *early* flies do not change their Ψ_CoM_, Ψ_Peak_, and R under any *T*-cycle, showing they are phase-locked to the lights-ON stimulus. Color codes: *early* - blue, *control* - green, *late* - orange.

Under T26, *early* flies showed similar emergence waveform as observed under T24 (*left panel third row, Fig. 4A).* The *control* flies showed phase advance *(mid panel third row, Fig. 4A)* and *late* flies showed even larger phase advance in response to T26 cycle *(right panel third row, Fig. 4A)* compared to their waveforms under T24. Although *early* flies showed higher percentages of “apparent” entrainment under the extreme *T*-cycles of T20 and T28, a very small fraction of *control* and *late* flies entrained. Thus, rendering comparison among stocks inappropriate and henceforth these results were not considered for further analyses. There is some degree of anticipation in *early* flies to lights-ON under all three *T*-cycles seen as gradual rise in emergence prior to lights-ON *(Fig. 4A).* This issue is addressed in *Supplementary Fig. S16.* Briefly, the underlying clock of *early* flies delay or advance under T22 and T26 respectively, still the high emergence immediately after lights-ON due to masking is conserved under all three *T*-cycles.

We quantified three parameters of the emergence waveform under all three *T*-cycles: a) Centre of Mass (Ψ_CoM_), which is an estimate of mean phase angle of emergence in a circular scale, b) Peak in ZT (Ψ_Peak_), and c) R, which is a measure of normalized amplitude of the eclosion rhythm and comprehensively describes consolidation of emergence. All these parameters have been previously used to describe the eclosion rhythm waveform of *Drosophila melanogaster* (Abhilash et al., 2019).

As seen in *Fig. 4B*, Ψ_CoM_ for *early* flies did not change across the *T*-cycles, whereas Ψ_CoM_ for *control* and *late* flies shifted in expected directions for T22 (delayed) and T26 (advanced) significantly compared to T24 cycle *(left panel, Fig. 4B; F*_4,12_ = 134.285, *p* < 0.05; *Supplementary table S4).* Ψ_Peak_ for *early* flies remained similar across all three *T*-cycles *(right panel, Fig. 4B; F*_4,12_ = 73.030,*p* < 0.05; *Supplementary table S5). control* and *late* flies showed expected trends in change of direction of peak phase shift in T22 and T26 cycles *(right panel, Fig. 4B; F*_4,12_ = 73.030, *p* < 0.05; *Supplementary table S5).* In case of *control* flies, although there was a trend of advancing peak phase in T26 cycle, compared to T24, this difference was not significant, whereas the delay in peak phase in T22 was much larger when compared to T24 cycle (*right panel, Fig. 4B; F*_4,12_ = 73.030,*p* < 0.05; *Supplementary table S5).* In *late* flies, the delay in peak phase in T26 was significantly larger when compared to T24 cycle, but the phase advance in T26 was not significant compared to T24 *(right panel, Fig. 4B; F*_4,12_ = 73.030, *p* < 0.05; *Supplementary table S5*).

Next, we quantified the consolidation of emergence/ normalized amplitude of the eclosion rhythm (R) of all populations under all *T*-cycles. A large R value is characteristic of the *early* population as evident by their narrow gate-width of emergence (Nikhil, Abhilash, et al., 2016). If the shorter and longer *T*-cycles had indeed shifted the circadian clock of *early* flies and the masking component is only responsible for the high emergence in the earliest part of the photophase, then value of R is expected to change among different *T*-cycles. Also, decrease in R may indicate one clock-controlled peak and one masking peak, as previously reported in case of locomotor activity rhythm in Drosophilid species (Prabhakaran and Sheeba, 2012; Rieger et al., 2003). We observed that R is significantly higher in *early* populations in all three *T*-cycles which suggests that they maintain constant high consolidation of emergence as observed previously under T24 (*middle panel, Fig. 4B; F*_4,12_ = 8.03, *p* < 0.05; *Supplementary table S6). control* flies do not change R among *T*-cycles, but *late* flies show significantly higher R under T26 compared to T24 and T22, primarily because of high amplitude of their emergence under T26.

Taken together, these results indicate that peak phase of *early* flies does not change when they entrain to short or long *T*-cycles. The fact that emergence occurs maximally at similar ZT across *T*-cycles supports the idea of larger masking component to this synchronization to *T*-cycles than a circadian clock mediated entrainment.

### *early* flies show phase lability under skeleton *T*-cycles

As we observed from previous experiments (*Fig. 2, 3, 4*), there is some degree of anticipation, a hallmark of circadian clock controlled rhythms, present under full *T*-cycles, we wanted to know to what extent circadian clock controls the phase of emergence in our stocks. However, it is masked by the high emergence immediately after lights-ON in *early* flies under full *T*-cycles. The nonparametric model of circadian entrainment posits that lights during the dawn and dusk transitions entrain the clock and even short light pulses at these phases are sufficient to reproduce the waveform of the rhythm seen under full photoperiod. Masking responses to light depend both on the duration of the illumination, and the intensity of light (Mrosovsky, 1999). We carried out skeleton photoperiod experiments to examine the extent of circadian clock control over the phase of emergence in our stocks, with short duration (0.25 hours) light pulses of ~70 lux and asked if these pulses elicit a masking response as well.

Although all populations showed entrainment under all skeleton *T*-cycles provided, their eclosion waveforms did not closely mimic those under full photoperiods of respective *T*-cycles *(Fig. 4A & 5A).* Under skeleton *T*-cycles, *early* flies became phase labile *(left and right panel, Fig. 5B).* With long *T*-cycles they advanced their phases, just as *control* and *late* flies *(Fig. 5A & 5B).* The change in Ψ_CoM_ across different *T*-cycles was small for *early* flies, compared to *control* and *late* flies *(left panel, Fig. 5B*).

**Figure 5:**
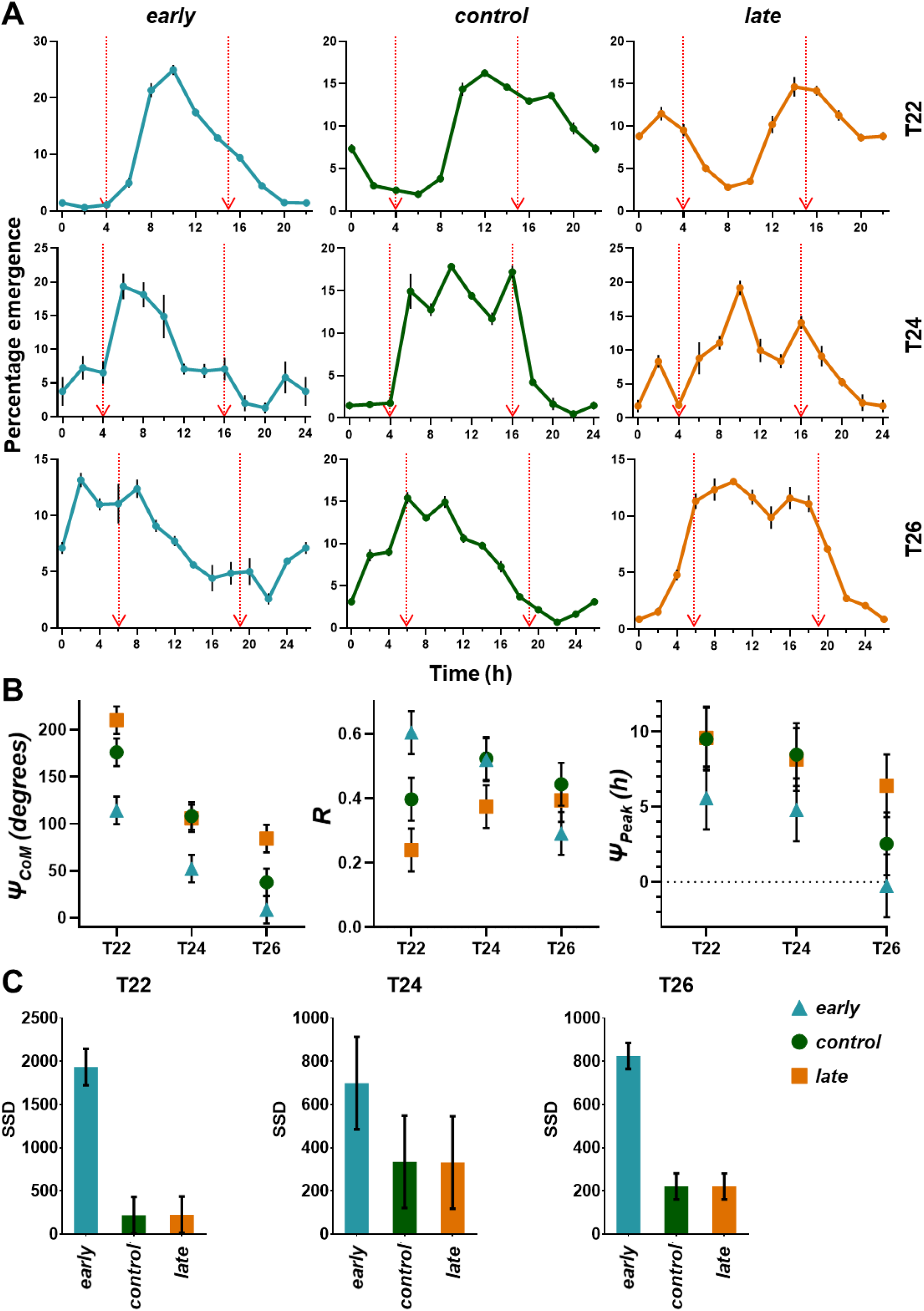
Emergence profile of *early*, *control*, and *late* flies in T22, T24, and T26 skeleton photoperiods and quantification of different eclosion rhythm parameters under different skeleton *T*-cycles. **A.** *early* flies change their emergence profile depending on skeleton *T*-cycles as do *control* and *late* flies. Red arrows depict the 15 minutes of the day when flies get light in respective skeleton photoperiods. Error bars are ± SEM. **B.** Left panel: Centre of Mass (Ψ_CoM_ - in degrees), middle panel: Consolidation of emergence (R), right panel: Peak in ZT (Ψ_Peak_, in hours). Error bars are ±95% CI. **C.** Sum of Square Difference (SSD) of waveforms under full and skeleton photoperiods. Significantly high SSD in *early* flies depict high amount of changes in waveform and phases between full and skeleton photoperiods. Error bars are ±95% CI.

Eclosion waveform of *early* flies showed large changes (in SSD) under skeleton *T*-cycles, whereas the changes were very small in case of *control* and *late* flies *(Fig. 5C; F*_2,6_ = 102.9092, *p* < 0.05; *F*_2,6_ = 158.3165, *p* < 0.05; *Supplementary tables S7, S9* - T22 and T26 respectively). We also compared the Ψ_CoM_ of the eclosion rhythm of all populations under different lengths of skeleton *T*-cycles. Ψ_Peak_ of the eclosion rhythm of all populations showed trends similar to Ψ_CoM_, but was not significantly different among stocks across *T*-cycles *(right panel, Fig. 5B; F*_4,12_ = 2.294, *p* > 0.05*; Supplementary table S12*) mostly due to the fact that the eclosion waveforms were not strictly unimodal in all individual vials. Although, *early* flies showed phase lability under skeleton *T*-cycles, they also showed a significantly advanced phase of entrainment (Ψ_CoM_) than *control* and *late* flies, across all three regimes *(bottom panel, Fig. 5B; F*_4,12_ = 7.462, *p* < 0.05; *Supplementary table S11*), suggesting that in addition to the masking response, advanced phase of circadian entrainment has also been selected for in our *early* flies. The extent of change in eclosion waveform between skeleton and full *T*-cycles can be seen in the change in R values (*Fig. 5B*). The *early* flies show significantly lower R value under T26 skeleton *T*-cycle, compared to the same under T22 and T24 skeleton *T*-cycles (*bottom panel, Fig. 5B; F*_4,12_ = 23.706, *p* < 0.05; *Supplementary table S10).*

These results suggest that, in *early* flies, although positive masking responses to lights-ON strongly influences Ψ_CoM_ and Ψ_Peak_ of the eclosion rhythm under full photoperiods, under skeleton photoperiods, these phases are clock controlled and significantly advanced (only Ψ_CoM_) compared to *control* and *late* flies.

## Discussion

While there have been several cases for the existence of masking as a phenomenon mediating biological rhythmicity (Aschoff, 1960b; Aschoff and von Goetz, 1988a; Binkley et al., 1983; Redlin and Mrosovsky, 1999), it has been long neglected by circadian biologists; Nicholas Mrosovsky (1999) wrote *“Nevertheless, as a phenomenon worth study in itself, masking generally has been neglected by circadian biologists. Their attention has been focused more on making sure that interpretations involving masking can be excluded in research on rhythms and on devising ways of eliminating masking effects from measurements of clock phase; these include the use of constant darkness and skeleton photoperiods”* (Mrosovsky, 1999).

For the most part, masking has received attention in study of locomotor activity rhythm (Fry, 1947; Sheeba et al., 2002), although recently it has been invoked to explain aspects of eclosion rhythm in *Drosophila* (McNabb and Truman, 2008; Thakurdas et al., 2009) and the silk moth *Bombyx mori* (Ikeda et al., 2019). It is known that the lights-ON signal mediated masking of *Drosophila* eclosion rhythm is brought about by the release of eclosion hormone (EH) as removal of EH neurons attenuates the masking response (McNabb and Truman, 2008). A sudden temperature change can also induce masking response in eclosion via the EH neurons (Jackson et al., 2005). There is indirect evidence that masking provides some adaptive value (Bloch et al., 2013; Lu et al., 2010), however to the best of our knowledge, there is no experimental evidence for masking evolving in response to selection.

### Evolution of masking in *early* populations along with advanced phase of entrainment

We hypothesized that due to the temporal placement of our selection window for the *early* flies (ZT22-ZT1), we may have inadvertently selected for individuals evolving a strong masking response to lights-ON (ZT0-ZT1), as we see in *Fig. 2A & 3A,* with very few (<5%) flies emerging in the window of ZT22-ZT0. Also, we observed 25-35% flies of *early* populations emerge immediately within half an hour after lights-ON (Ψ_Peak_ = ZT0.5) *(Fig. 2A & 3A),* which led us to think there might be a significant masking component regulating the phase of the eclosion rhythm of *early* flies.

Unlike *control* and *late* populations, upon advancement of lights-ON, many of the *early* flies eclose immediately after lights-ON rather than what we would have expected if their eclosion rhythm was strongly circadian clock driven *(Fig. 2A & 2B).* Alternatively, this result suggests that the light-sensitive A-oscillator in *early* flies is stronger or dominant and facilitates re-entrainment in the very next cycle. However, as discussed below, the lights-ON response in eclosion is not as immediate as for other behaviors, such as locomotor activity rhythm, so the high emergence immediately after lights-ON on *Assay day 2 (Fig. 2A*) being a result of faster re-entrainment seems highly unlikely. Our hypothesis of strong masking for the *early* flies was further validated by lower emergence in *early* population at subjective dawn in DD *(Fig. 3A & 3B)* in contrast to the flies from the other two sets of populations. This result may also be interpreted as - *early* flies showing higher amplitude expansion under entrainment for eclosion rhythms followed by amplitude reduction in response to transition from LD to DD. However, the high emergence at ZT0.5 (presumably the masking induced peak) under LD is absent in DD, whereas the next highest peak at ZT1 (presumably the clock-controlled peak) under LD is maintained at similar phase under DD (~10% emergence), showing a phase control and is seen in previous experiments as well (*Fig. 2A*). While *control* and *late* flies phased eclosion in the expected direction based on circadian clock control under short and long *T*-cycles *(Fig. 4A & 5A),* the *early* flies consistently respond by positive masking with high emergence immediately after lights-ON. However, the response of *early* flies under skeleton pulse-induced *T*-cycles was remarkable because similar to the *control* and *late* flies, they too delayed and advanced emergence phases (Ψ_CoM_) under T22 and T26 respectively (*Fig. 5*).

In the phase advance experiments (*Fig. 2*), although *early* flies exhibited high emergence immediately after lights-ON at ZT0.5, there was considerable emergence even around ZT1.5 - 2 - this peak was comparable to the peak at ZT1 on *Assay day 2* of LD-DD transition experiments (*bottom panel, Fig. 2A & bottom panel, Fig. 3A*). We reasoned that the phase advance of lights-ON on *Assay day 2* exposed the late night (advance zone) of the PRC of *early* flies to light (Kumar et al., 2007), thus advancing the clock and hence producing an advanced peak (at ZT 2 in phase advance experiments and ZT1 in LD-DD transition experiments - both at *Assay day 2*). The exposure to three skeleton *T*-cycles revealed that indeed *early* flies maintained significantly advanced Ψ_CoM_ compared to *control* and *late* flies *(Fig. 5B),* suggesting that our selection protocol has indeed resulted in advanced phase of emergence in *early* flies along with greater masking which is revealed under full photoperiods. The fact that the *control* flies show some extent of masking (*Fig. 3B*), and upon selection for advanced phase of entrainment, can give rise to significantly higher masking (*early* population), suggests that masking-inducing variations are present in populations, albeit in low frequency. Recently, Pegoraro and colleagues have shown that negative masking can evolve when flies are selected for nocturnality (Pegoraro et al., 2019).

### Higher range of “apparent” entrainment achieved by masking

In another set of experiments in which all populations were subjected to T20 and T28 cycles, *early* flies showed much higher percentages of apparent entrainment compared to *control* and *late* flies (*Supplementary fig. S15*). This suggested that, though *early* flies show entrainment, whereas *control* and *late* flies show almost none, *early* flies may mask successfully and show an apparent entrained behavior. This shows that a spectacularly high range of “apparent entrainment” can be achieved just by masking, at least for the eclosion rhythm. Altogether our experiments make a very strong case that in the process of creating the *early* chronotypes, in addition to them evolving a faster clock with advanced phase of entrainment, they have also evolved robust positive masking responses to lights-ON. This facilitates the idea that in nature, organisms may use masking - a clock-independent phenomenon, as a mechanism to phase themselves to appropriate times of the day, e.g., eclosion being gated to the early part of the day to prevent desiccation and enhance survival rate. It can be argued that masking maybe an evolutionary disadvantage in the sense that to compensate for a highly “noisy” clock masking may evolve to maintain specific phases locked to a zeitgeber(s) and does not have the flexibility of phase lability in complex environments. Indeed, previous work from our lab has shown under complex zeitgeber conditions (light:dark 12:12 + warm:cool 12:12; in-phase and out-of-phase), *early* flies show remarkable resilience to shift phases and stay phase locked to the lights-ON signal, whereas *control* and *late* flies change their phases more readily (Abhilash et al., 2019). However, empirical evidence suggests that a fully functional clock and masking can co-exist in an animal and in all probability these are complementary mechanisms for organisms to maintain specific phases (Aschoff and von Goetz, 1988a; Mrosovsky, 1994; Redlin and Mrosovsky, 1999, 2004; Rensing, 1989).

### The duration and intensity of light dramatically alters magnitude of response in eclosion

The light used in all our experiments was of ~70 lux (to match the long term selection maintenance regime), and can be considered low intensity compared to majority of the eclosion rhythm studies in *Drosophila* (which typically used ~750 lux). It is especially so in those studies where relationship between the lights-ON signal and downstream pathways have been analyzed and linked to timing of eclosion (Baker et al., 1999; McNabb and Truman, 2008). They show that lights-ON signal can rapidly induce eclosion of up to ~20% of the waveform within about 10 minutes. This suggests that a skeleton photoperiod should be able to induce high masking responses, which we did not observe in any of the skeleton *T*-cycles. This can be explained by the different light intensity used in our experiments (at least 10 times lower). Nevertheless, our regime induces ~35% emergence in a mere 30 minutes when the lights-ON was advanced *(Fig. 2B).* We propose that similar number of photons integrated over time could induce high emergence in *early* populations, compared to *control* and *late* populations.

Another important difference from previous studies which were mostly aimed to understand the mechanisms of EH release and downstream pathways leading to the act of eclosion is, that they used cultures containing similar developmental stages. To observe light-induced emergence, typically experiments were set up such that flies were expected to emerge in the next ~60 minutes after light pulses were given (McNabb and Truman, 2008). Our studies together with this information from previous reports lead us to hypothesize that photon integration is a part of this masking response, and that as soon as a threshold is crossed, masking response to lights-ON is observed.

### Possible mechanisms driving higher masking in *early* populations

Some hint of the evolution of masking mechanisms are seen in an *in-silico* study, where gene regulatory networks were allowed to evolve under selection for correct prediction of phases under light-dark cycles (Troein et al., 2009). It was seen that under a simple LD12:12 condition, only delayed light responses were selected for, while no oscillatory mechanism was found to evolve. This suggests that in a minimal environment, where the only cycling cue is a light-dark cycle, oscillators may not be necessary to achieve particular phases, and a delayed light response may as well do the trick. In our selection regime, environmental conditions are similarly minimal, with only one cycling cue, a ~70 lux LD12:12 square light cycle (abrupt transitions) regime with constant ambient temperature of 25±0.5 °C, and results from Troein and colleagues supports the idea of evolution of simpler phasing mechanisms, such as strong masking.

This evolved masking response may be governed by co-evolved non-circadian photosensitivity. Unpublished work from our laboratory shows that rhodopsins *(Rh4, Rh7)* and non-rhodopsin photosensitive molecules like *ninaC (neither inactivation nor afterpotential C* - known to affect phototransduction and responses to light stimulus) have accumulated more SNPs in genome of *early* flies when compared to *control* genomes (*manuscript under preparation, Abhilash Lakshman, Arijit Ghosh, K L Nikhil).* We also find that *early* flies have accumulated more SNPs in genes implicated in eye physiology (*ruby*, *cardinal*, *carmine*, *scabrous*, and *friend of echinoid*) and that they show significantly high allele diversity compared to *late* flies in eclosion-related genes (*eyes absent, Ecdysone receptor, scabrous)* and circadian rhythm related genes *(Pigment-dispersing factor, Pigment-dispersing factor receptor, Ecdysone receptor, shaggy) (manuscript under preparation, Abhilash Lakshman, Arijit Ghosh, K L Nikhil).* Thus, there are the possibilities of (a) strong photosensitivity and EH neurons coupling or (b) heightened photosensitivity of the PG itself having evolved in the *early* flies.

To the best of our knowledge this work is the first experimental demonstration that masking can evolve as a response to selection for phase of entrainment. We propose that masking can be a valid mechanism by which organisms show *early* chronotype in an environment where strong light transitions are present. Further investigations into the molecular mechanisms and neuronal control of this masking response is needed, for which some plausible targets have been mentioned above. Overall, this work highlights the complex mechanisms of “apparent” light entrainment and provides an experimental framework to dissect out relative contributions of masking and the circadian clock regulating timing of a behavior.

## Supporting information

All supplementary data

## Acknowledgements

We are extremely grateful to late Professor Vijay Kumar Sharma for providing us with this beautiful model system to address such questions regarding phase of entrainment of the circadian clock. The maintenance of these populations throughout last ~17 years have seen multiple graduate and masters students working relentlessly day and night, and we stand on the shoulders of the vast knowledge created by the previous members of the lab and take it forward. The assays carried out in this body of work included presence and intense work for 24 hours at a stretch of 4-5 days, which was not humanely possible for a handful of students, and we acknowledge help of summer interns (Sankeert Satheesan and Bharathy Nagarajan) in initial experiments of standardization. We thank Dr. Abhilash Lakshman for carefully reading our manuscript and suggesting some very useful changes. We would also like to acknowledge financial support from the Science and Engineering Research Board, New Delhi, to V.S. (CRG/2019/006802); intramural funding from Jawaharlal Nehru Centre for Advanced Scientific Research (JNCASR), Bangalore; the Council of Scientific & Industrial Research (CSIR), New Delhi, India, for a junior research fellowship and a senior research fellowship to A.G.; JNCASR, for a fellowship to P.S; the Department of Science and Technology, New Delhi, India for an INSPIRE fellowship for masters studies to SD; and consumable grant from the Department of Biotechnology (DBT), Government of India, to V.S. (LSRET-JNC/SV/4532/19-20).

