## Supplementary material for "Evidence for co-evolution of masking and circadian phase in *Drosophila melanogaster*": All supplementary data

*Running title: To mask or not to mask?*

*Correspondence: Vasu Sheeba, Chronobiology and Behavioral Neurogenetics Laboratories, Neuroscience Unit, Jawaharlal Nehru Centre for Advanced Scientific Research, Jakkur, Bengaluru - 560064, Karnataka, India. Phone: +91-080-2208-2987.

**Statistically significant fixed factors or interaction terms are *italicized* in all ANOVA tables.**

**Supplementary Table S1: ANOVA table summarizing the effect of Selection on the difference in percentage emergence between pre- and post-masking days in the window specified in Figure. 3 (Dashed rectangle).**

| | **Effect (F/R)** | **SS** | **Df** | **MS** | **Error df** | **Error MS** | **F** | **p** |
|---|---|---|---|---|---|---|---|---|
| ***Intercept*** | *Fixed* | *144.0970* | *1* | *144.0970* | *3* | *0.368485* | *391.0525* | *0.000283* |
| ***Selection*** | *Fixed* | *248.8150* | *2* | *124.4075* | *6* | *0.621080* | *200.3084* | *03* |
| **Block** | Random | 1.1055 | 3 | 0.3685 | 6 | 0.621080 | 0.5933 | 0.641946 |
| **Selection*Block** | Random | 3.7265 | 6 | 0.6211 | 0 | 0 | | |

**Supplementary Table S2: ANOVA table summarizing the effect of Selection on the difference in percentage emergence between pre- and post-masking days in the window specified in Figure. 3 (Solid rectangle).**

| | **Effect (F/R)** | **SS** | **Df** | **MS** | **Error df** | **Error MS** | **F** | **p** |
|---|---|---|---|---|---|---|---|---|
| ***Intercept*** | *Fixed* | *1660.226* | *1* | *1660.226* | *3* | *6.69329* | *248.0432* | *0.000556* |
| ***Selection*** | *Fixed* | *1485.424* | *2* | *742.712* | *6* | *16.07567* | *46.2010* | *0.000227* |
| **Block** | Random | 20.080 | 3 | 6.693 | 6 | 16.07567 | 0.4164 | 0.747797 |
| **Selection*Block** | Random | 96.454 | 6 | 16.076 | 0 | 0 | | |

**Supplementary Table S3: ANOVA table summarizing the effect of Selection on the difference of percentage emergence between pre-masking and post-masking days from ZT0-0.5.**

| | **Effect (F/R)** | **SS** | **Df** | **MS** | **Error df** | **Error MS** | **F** | **p** |
|---|---|---|---|---|---|---|---|---|
| ***Intercept*** | *Fixed* | *1123.801* | *1* | *1123.801* | *3* | *8.284387* | *135.6528* | *0.001360* |
| ***Selection*** | *Fixed* | *684.151* | *2* | *342.075* | *6* | *6.171058* | *55.4322* | *0.000135* |
| **Block** | Random | 24.853 | 3 | 8.284 | 6 | 6.171058 | 1.3425 | 0.346128 |
| **Selection*Block** | Random | 37.026 | 6 | 6.171 | 0 | 0 | | |

**Supplementary Table S4: ANOVA table summarizing the effects of Selection, full *T*-cycle and their interactions on $\Psi_{CoM}$.**

| | **Effect (F/R)** | **SS** | **Df** | **MS** | **Error df** | **Error MS** | **F** | **p** |
|---|---|---|---|---|---|---|---|---|
| ***Intercept*** | *Fixed* | *311448.8* | *1* | *311448.8* | *3* | *57.97748* | *5371.892* | *06* |
| ***Selection*** | *Fixed* | *57207.8* | *2* | *28603.9* | *6* | *87.12321* | *328.316* | *01* |
| ***T-cycle*** | *Fixed* | *49298.1* | *2* | *24649.0* | *6* | *37.50927* | *657.145* | *0* |
| **Block** | Random | 173.9 | 3 | 58.0 | 4.68156 | 87.01872 | 0.666 | 0.609976 |

| | | | | | | | | |
|---|---|---|---|---|---|---|---|---|
| ***Selection*T-cycle*** | *Fixed* | *20203.8* | *4* | *5051.0* | *12* | *37.61376* | *134.285* | *0* |
| **Selection*Block** | Random | 522.7 | 6 | 87.1 | 12 | 37.61376 | 2.316 | 0.101623 |
| **T-cycle*Block** | Random | 225.1 | 6 | 37.5 | 12 | 37.61376 | 0.997 | 0.469741 |
| **Selection*T-cycle*Block** | Random | 451.4 | 12 | 37.6 | 0 | 0 | | |

**Supplementary Table S5: ANOVA table summarizing the effects of Selection, full *T*-cycle and their interactions on $\Psi_{Peak}$.**

| | Effect (F/R) | SS | Df | MS | Error df | Error MS | F | p |
|---|---|---|---|---|---|---|---|---|
| ***Intercept*** | *Fixed* | *959.2441* | *1* | *959.2441* | *3* | *0.505395* | *1898.007* | *0.000027* |
| ***Selection*** | *Fixed* | *283.4216* | *2* | *141.7108* | *6* | *1.010729* | *140.207* | *09* |
| ***T-cycle*** | *Fixed* | *201.8840* | *2* | *100.9420* | *6* | *0.313529* | *321.955* | *01* |
| **Block** | Random | 1.5162 | 3 | 0.5054 | 3.41087 | 0.838812 | 0.603 | 0.651883 |
| ***Selection*T-cycle*** | *Fixed* | *141.8077* | *4* | *35.4519* | *12* | *0.485445* | *73.030* | *0* |
| **Selection*Block** | Random | 6.0644 | 6 | 1.0107 | 12 | 0.485445 | 2.082 | 0.131734 |
| **T-cycle*Block** | Random | 1.8812 | 6 | 0.3135 | 12 | 0.485445 | 0.646 | 0.693179 |
| **Selection*T-cycle*Block** | Random | 5.8253 | 12 | 0.4854 | 0 | 0 | | |

**Supplementary Table S6: ANOVA table summarizing the effects of Selection, full *T*-cycle and their interactions on R.**

| | Effect (F/R) | SS | Df | MS | Error df | Error MS | F | p |
|---|---|---|---|---|---|---|---|---|
| ***Intercept*** | *Fixed* | *13.95465* | *1* | *13.95465* | *3* | *0.000579* | *24096.85* | *01* |
| ***Selection*** | *Fixed* | *0.37793* | *2* | *0.18897* | *6* | *0.002806* | *67.34* | *0.000078* |
| **T-cycle** | Fixed | 0.00535 | 2 | 0.00267 | 6 | 0.001977 | 1.35 | 0.327323 |
| **Block** | Random | 0.00174 | 3 | 0.00058 | 8.79582 | 0.004187 | 0.14 | 0.934538 |
| ***Selection*T-cycle*** | *Fixed* | *0.01913* | *4* | *0.00478* | *12* | *0.000596* | *8.03* | *0.002174* |
| **Selection*Block** | Random | 0.01684 | 6 | 0.00281 | 12 | 0.000596 | 4.71 | 0.010906 |
| **T-cycle*Block** | Random | 0.01186 | 6 | 0.00198 | 12 | 0.000596 | 3.32 | 0.036548 |
| **Selection*T-cycle*Block** | Random | 0.00715 | 12 | 0.00060 | 0 | 0 | | |

**Supplementary Table S7: ANOVA table summarizing the effect of Selection on SSD calculated between T22 skeleton and full *T*-cycle profiles.**

| | **Effect (F/R)** | **SS** | **Df** | **MS** | **Error df** | **Error MS** | **F** | **p** |
|---|---|---|---|---|---|---|---|---|
| ***Intercept*** | *Fixed* | *7532872* | *1* | *7532872* | *3* | *9269.89* | *812.6171* | *0.000095* |
| ***Selection*** | *Fixed* | *7823911* | *2* | *3911955* | *6* | *38013.65* | *102.9092* | *0.000023* |
| **Block** | Random | 27810 | 3 | 9270 | 6 | 38013.65 | 0.2439 | 0.862905 |
| **Selection*Block** | Random | 228082 | 6 | 38014 | 0 | 0.00 | | |

**Supplementary Table S8: ANOVA table summarizing the effect of Selection on SSD calculated between T24 skeleton and full *T*-cycle profiles.**

| | **Effect (F/R)** | **SS** | **Df** | **MS** | **Error df** | **Error MS** | **F** | **p** |
|---|---|---|---|---|---|---|---|---|
| ***Intercept*** | *Fixed* | *2484756* | *1* | *2484756* | *3* | *13279.91* | *187.1063* | *0.000845* |
| **Selection** | Fixed | 356923 | 2 | 178461 | 6 | 38904.85 | 4.5871 | 0.061821 |
| **Block** | Random | 39840 | 3 | 13280 | 6 | 38904.85 | 0.3413 | 0.796850 |
| **Selection*Block** | Random | 233429 | 6 | 38905 | 0 | 0.00 | | |

**Supplementary Table S9: ANOVA table summarizing the effect of Selection on SSD calculated between T26 skeleton and full *T*-cycle profiles.**

| | **Effect (F/R)** | **SS** | **Df** | **MS** | **Error df** | **Error MS** | **F** | **p** |
|---|---|---|---|---|---|---|---|---|
| ***Intercept*** | *Fixed* | *2137588* | *1* | *2137588* | *3* | *7183.089* | *297.5862* | *0.000424* |
| ***Selection*** | *Fixed* | *973610* | *2* | *486805* | *6* | *3074.883* | *158.3165* | *06* |
| **Block** | Random | 21549 | 3 | 7183 | 6 | 3074.883 | 2.3361 | 0.173219 |
| **Selection*Block** | Random | 18449 | 6 | 3075 | 0 | 0.000 | | |

**Supplementary Table S10: ANOVA table summarizing the effects of Selection, skeleton *T*-cycle and their interactions on R.**

| | **Effect (F/R)** | **SS** | **Df** | **MS** | **Error df** | **Error MS** | **F** | **p** |
|---|---|---|---|---|---|---|---|---|
| ***Intercept*** | *Fixed* | *6.366578* | *1* | *6.366578* | *3* | *0.003187* | *1997.577* | *0.000025* |
| ***Selection*** | *Fixed* | *0.131338* | *2* | *0.065669* | *6* | *0.001436* | *45.729* | *0.000233* |
| ***T-cycle*** | *Fixed* | *0.056974* | *2* | *0.028487* | *6* | *0.003262* | *8.733* | *0.016717* |
| **Block** | Random | 0.009561 | 3 | 0.003187 | 1.72617 | 0.002143 | 1.487 | 0.446379 |
| ***Selection*T-cycle*** | *Fixed* | *0.242255* | *4* | *0.060564* | *12* | *0.002555* | *23.706* | *0.000013* |
| **Selection*Block** | Random | 0.008616 | 6 | 0.001436 | 12 | 0.002555 | 0.562 | 0.752789 |
| **T-cycle*Block** | Random | 0.019572 | 6 | 0.003262 | 12 | 0.002555 | 1.277 | 0.337396 |

| | | | | | | | | |
|---|---|---|---|---|---|---|---|---|
| **Selection*T-cycle*Block** | Random | 6.366578 | 1 | 6.366578 | 3 | 0.003187 | 1997.577 | 0.000025 |

**Supplementary Table S11: ANOVA table summarizing the effects of Selection, skeleton *T*-cycle and their interactions on $\Psi_{CoM}$.**

| | Effect (F/R) | SS | Df | MS | Error df | Error MS | F | p |
|---|---|---|---|---|---|---|---|---|
| ***Intercept*** | *Fixed* | *358228.4* | *1* | *358228.4* | *3* | *263.8468* | *1357.714* | *0.000044* |
| ***Selection*** | *Fixed* | *34817.6* | *2* | *17408.8* | *6* | *340.8811* | *51.070* | *0.000171* |
| ***T-cycle*** | *Fixed* | *93361.1* | *2* | *46680.5* | *6* | *106.9907* | *436.305* | *0* |
| **Block** | Random | 791.5 | 3 | 263.8 | 4.63149 | 323.2988 | 0.816 | 0.541237 |
| ***Selection*T-cycle*** | *Fixed* | *3718.3* | *4* | *929.6* | *12* | *124.5731* | *7.462* | *0.002935* |
| **Selection*Block** | Random | 2045.3 | 6 | 340.9 | 12 | 124.5731 | 2.736 | 0.065054 |
| **T-cycle*Block** | Random | 641.9 | 6 | 107.0 | 12 | 124.5731 | 0.859 | 0.550706 |
| **Selection*T-cycle*Block** | Random | 1494.9 | 12 | 124.6 | 0 | 0.0000 | | |

**Supplementary Table S12: ANOVA table summarizing the effects of Selection, skeleton *T*-cycle and their interactions on $\Psi_{Peak}$.**

| | Effect (F/R) | SS | Df | MS | Error df | Error MS | F | p |
|---|---|---|---|---|---|---|---|---|
| ***Intercept*** | *Fixed* | *1336.309* | *1* | *1336.309* | *3* | *0.348148* | *3838.333* | *09* |
| ***Selection*** | *Fixed* | *140.842* | *2* | *70.421* | *6* | *2.664815* | *26.426* | *0.001060* |
| ***T-cycle*** | *Fixed* | *190.582* | *2* | *95.291* | *6* | *2.854815* | *33.379* | *0.000561* |
| **Block** | Random | 1.044 | 3 | 0.348 | 2.91984 | 2.995370 | 0.116 | 0.944653 |
| **Selection*T-cycle** | Fixed | 23.166 | 4 | 5.791 | 12 | 2.524259 | 2.294 | 0.119177 |
| **Selection*Block** | Random | 15.989 | 6 | 2.665 | 12 | 2.524259 | 1.056 | 0.438650 |
| **T-cycle*Block** | Random | 17.129 | 6 | 2.855 | 12 | 2.524259 | 1.131 | 0.401340 |
| **Selection*T-cycle*Block** | Random | 30.291 | 12 | 2.524 | 0 | 0 | | |

**Supplementary Table S13: $\Psi_{CoM}$, $\Psi_{Peak}$, R, SSD values of *early*, *control* and *late* populations under full or skeleton *T*-cycles and between them.**

| | | **Full *T*-cycles** | | | **Skeleton *T*-cycles** | | | **SSD values** | | | |
|---|---|---|---|---|---|---|---|---|---|---|---|
| | | **T22** | **T24** | **T26** | **T22** | **T24** | **T26** | | **T22** | **T24** | **T26** |
| ***early*** | **$\Psi_{CoM}$** | 53.3342 | 49.3746 | 39.9952 | 114.275 | 52.3852 | 8.6341 | ***early*** | 1934.22 | 698.935 | 824.883 |
| | **$\Psi_{Peak}$** | 1.95 | 2.05 | 2 | 5.6 | 4.8 | -0.25 | ***control*** | 218.942 | 334.61 | 220.793 |
| | **R** | 0.75923 | 0.76716 | 0.74359 | 0.60375 | 0.51946 | 0.29064 | ***late*** | 223.74 | 331.58 | 220.496 |
| ***control*** | **$\Psi_{CoM}$** | 141.691 | 83.2699 | 35.5682 | 176.087 | 108.232 | 37.7991 | | | | |
| | **$\Psi_{Peak}$** | 9.65 | 2.8 | 1.55 | 9.5 | 8.48 | 2.55 | | | | |
| | **R** | 0.61622 | 0.58808 | 0.60528 | 0.39672 | 0.52365 | 0.44359 | | | | |
| ***late*** | **$\Psi_{CoM}$** | 219.701 | 146.947 | 67.2327 | 210.211 | 105.923 | 84.2354 | | | | |
| | **$\Psi_{Peak}$** | 12.66 | 10.4875 | 3.31 | 9.6 | 8.15 | 6.4 | | | | |
| | **R** | 0.48222 | 0.47371 | 0.56791 | 0.23944 | 0.37427 | 0.39324 | | | | |

Note: All phase values are phase relationships with lights-ON.

**Supplementary Figure S14: Difference in percentage emergence between pre-masking and post-masking days in two 2.5-hour time windows.** The left panels show time windows used for analysis (red and black rectangles respectively). In same time windows *early* flies show significantly higher (red region) or lower (black region) emergence than that of *control* and *late* flies on post-masking day than pre-masking day, showing this high emergence immediately after lights-ON is specific to *early* flies.

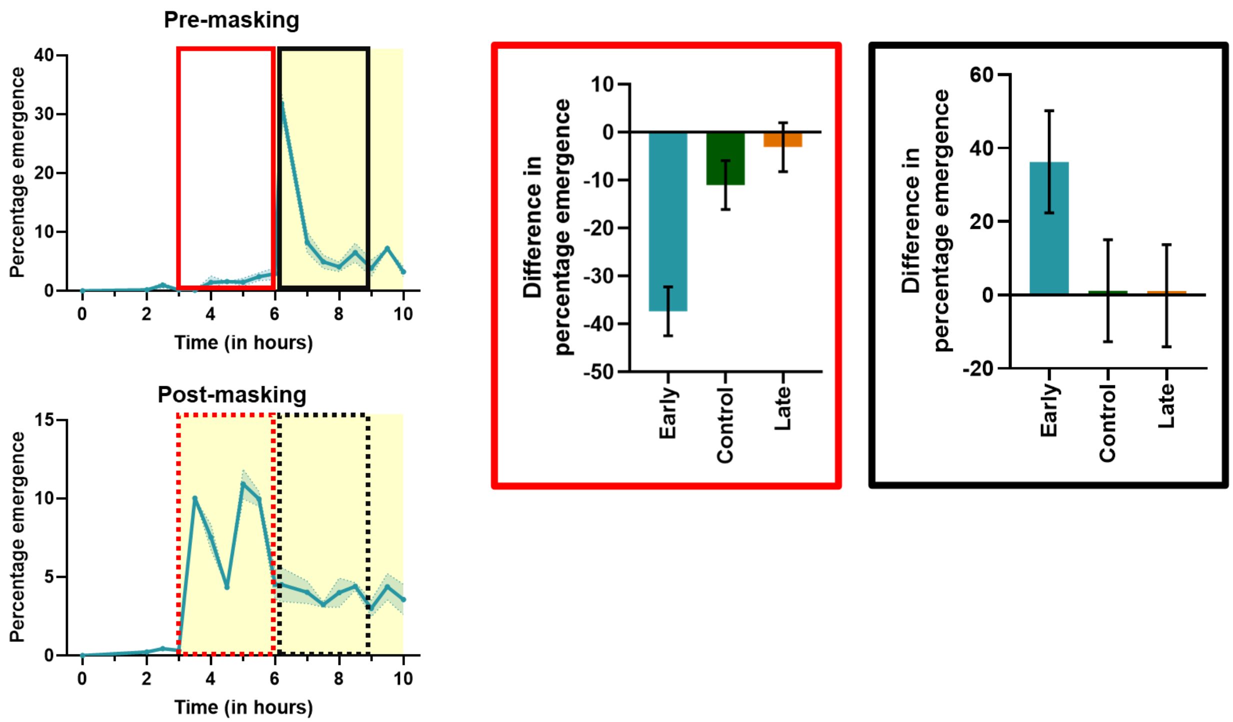


**Supplementary Figure S15: Percentage entrainment of *early*, *control* and *late* flies under T20 and T28.**

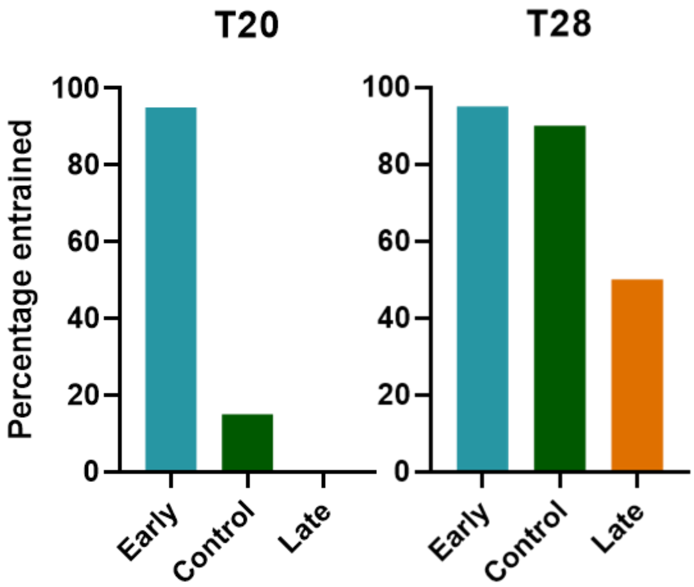


**Supplementary Figure S16: Percentage emergence of *early* flies under full *T*-cycles (T22, T24 and T26) 2, 4, and 6 hours before lights-ON.** As postulated in the main text, the putative clock-controlled peak for *early* flies occurs at ZT1 under LD12:12 (*Fig. 2A* and *3A*). The anticipation to this clock-controlled peak can be seen under all *T*-cycles in form of emergence before lights-ON (*Fig. 4A, left column*). Though the masking-induced high emergence immediately after lights-ON at ZT2 is conserved under all three *T*-cycles (*Fig. 4A, left column*), the emergence starts much earlier under T26 (compared to under T22 and T24) as evident by significantly high emergence 6 hours before lights-ON, as though the underlying clock has advanced. On the other hand, under T22, the emergence 2 hours prior to lights-ON is similar to T24 and T26, while it is almost zero prior to this, as if the underlying clock is delayed compared to the other *T*-cycles. Error bars are ±95% CI (main effect of *T-cycle*, three separate ANOVAs for three different time points – *Supplementary table S17, S18 and S19*).

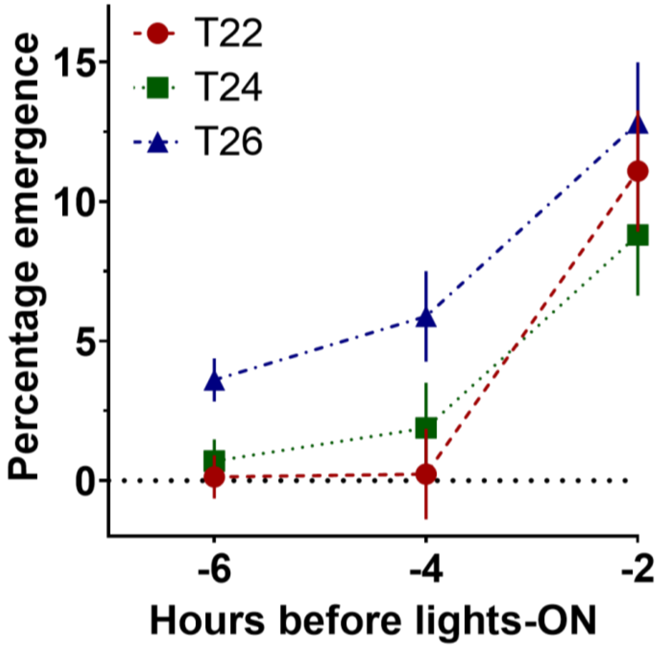


**Supplementary Table S17: ANOVA table summarizing the effect of *T*-cycle on percentage emergence at 2 hours prior to lights-ON under three full *T*-cycles in early flies.**

| | Effect (F/R) | SS | Df | MS | Error df | Error MS | F | p |
|---|---|---|---|---|---|---|---|---|
| ***Intercept*** | *Fixed* | *1426.558* | *1* | *1426.558* | *3* | *8.240409* | *173.1174* | *0.000948* |
| **T-cycle** | Fixed | 32.386 | 2 | 16.193 | 6 | 4.020893 | 4.0272 | 0.077807 |
| **Block** | Random | 24.721 | 3 | 8.240 | 6 | 4.020893 | 2.0494 | 0.208542 |
| **T-cycle*Block** | Random | 24.125 | 6 | 4.021 | 0 | 0 | | |

**Supplementary Table S18: ANOVA table summarizing the effect of *T*-cycle on percentage emergence at 4 hours prior to lights-ON under three full *T*-cycles in early flies.**

| | Effect (F/R) | SS | Df | MS | Error df | Error MS | F | p |
|---|---|---|---|---|---|---|---|---|
| ***Intercept*** | *Fixed* | *85.29175* | *1* | *85.29175* | *3* | *2.082540* | *40.95564* | *0.007728* |
| ***T-cycle*** | *Fixed* | *67.70220* | *2* | *33.85110* | *6* | *2.239383* | *15.11626* | *0.004541* |
| **Block** | Random | 6.24762 | 3 | 2.08254 | 6 | 2.239383 | 0.92996 | 0.481873 |
| **T-cycle*Block** | Random | 13.43630 | 6 | 2.23938 | 0 | 0 | | |

**Supplementary Table S19: ANOVA table summarizing the effect of *T*-cycle on percentage emergence at 6 hours prior to lights-ON under three full *T*-cycles in early flies.**

| | Effect (F/R) | SS | Df | MS | Error df | Error MS | F | p |
|---|---|---|---|---|---|---|---|---|
| ***Intercept*** | *Fixed* | *26.25740* | *1* | *26.25740* | *3* | *0.262412* | *100.0618* | *0.002126* |
| ***T-cycle*** | *Fixed* | *27.85751* | *2* | *13.92876* | *6* | *0.511296* | *27.2421* | *0.000976* |
| **Block** | Random | 0.78724 | 3 | 0.26241 | 6 | 0.511296 | 0.5132 | 0.687986 |
| **T-cycle*Block** | Random | 3.06777 | 6 | 0.51130 | 0 | 0 | | |
